# The Roles of Different Spatial Frequency Channels in Real-World Visual Motion Perception

**DOI:** 10.1101/328443

**Authors:** Cong Shi, Shrinivas Pundlik, Gang Luo

## Abstract

Speed perception is an important task performed by our visual system in various daily life tasks. In various psychophysical tests, relationship between spatial frequency, temporal frequency, and speed has been examined in human subjects. The role of vision impairment in speed perception has also been previously examined. In this work, we examine the inter-relationship between speed, spatial frequency, low vision conditions, and the type of input motion stimuli in motion perception accuracy. For this purpose, we propose a computational model for speed perception and evaluate it in custom generated natural and stochastic sequences by simulating low-vision conditions (low pass filtering at different cutoff frequencies) as well as complementary vision conditions (high pass versions at the same cutoff frequencies). Our results show that low frequency components are critical for accurate speed perception, whereas high frequencies do not play any important role in speed estimation. Since perception of low frequencies may not be impaired in visual acuity loss, speed perception was not found to be impaired in low vision conditions compared to normal vision condition. We also report significant differences between natural and stochastic stimuli, notably an increase in speed estimation error when using stochastic stimuli compared to natural sequences, emphasizing the use of natural stimuli when performing future psychophysical studies for speed perception.

## Introduction

Visual motion perception plays an important role a wide range of tasks in an organism’s life, such as navigation or evading predators. The neural circuitry for motion perception can differ depending on the requirements. For example, simple but fast motion processing is important for a fly to evade a predator.(Borst and Euler, 2011) On the other hand, primates have the ability to process more complex motion information.(Orchard and Etienne-Cummings, 2014) Speed estimation is an important aspect of primate visual motion perception. Neurons in the MT region, as well as some complex V1 neurons are specially tuned for speed perception, (Perrone and Thiele, 2001; Simoncelli and Heeger, 2001 Liu and Newsome, 2002; Priebe et al., 2006) and lesions in MT area can lead to motion perception deficits.(Newsome and Pare, 1988) Particularly intriguing aspect of these speed tuned neurons in the primate MT region is that they are somewhat invariant to the spatial frequency of the stimuli, allowing for finer speed estimation. This is especially important given the wide band of spatiotemporal frequencies received by our visual system.

Speed estimation in visual motion depends upon the spatial and temporal frequency of the stimulus. In a simple example involving horizontally moving sinewave grating shown in Figure 4, the speed of motion is the ratio of temporal frequency and the spatial frequency. A speed tuned neuron thus has an oriented receptive field stretching over a range of spatial and temporal frequencies, with the slope defining the speed at which the neuron with fire.(Perrone and Thiele, 2001; Simoncelli and Heeger, 2001) Higher speed leads to an increase in the slope of the orientation of the receptive field, which means that the range of spatial frequencies at which higher speeds can be perceived is narrowed, with a shift towards lower frequency band (Ref. fig 6 in Perrone and Thiele). This phenomenon is seen repeatedly in a large number of psychophysical studies that measured motion perception thresholds, indicating a strong bias of the visual system toward low frequencies when perceiving higher speed motion. In other words, we can hypothesize that low frequencies are important in perception of higher speed motion. Studies involving blurred stimuli (no high frequency components) or people with low vision (people with reduced visual acuity that impairs their ability to perceive high frequency components) provide further indication of the importance of low frequency channels in motion perception.

Motion is a strong cue that allows extraction of valuable scene information, even when other visual cues corresponding to higher frequency components are not available: normally sighted subjects could not recognize events in natural scenes when presented with randomly picked highly blurred frames, but were able to identify events when presented as a continuous video stream.(Pan and Bingham, 2013) Similarly, subjects fitted with blur glasses simulating low visual acuity were able to correctly determine contents of video clips, just like those viewing contents without blur glasses.(Saunders et al., 2014) Studies measuring motion perception threshold using artificial stimuli showed that low vision only degraded the capability to perceive very slow motions (below 2 cycles/degree), and there was no significant difference in perception of faster motion in individuals with small to moderate VA loss compared to normally sighted subjects.(Yang and Stevenson, 1997; Lappin et al., 2009) Thus, a relationship exists between the spatial frequency channels, visual motion perception, and the visual acuity with an underlying hypothesis that the higher spatial frequency components are not critical as the lower frequency channels in motion perception when dealing with real-world scenarios.

In spite of multiple studies that support the above hypothesis, there are no compelling quantitative analyses of the roles of different spatial frequency channels in perception of motion speed in natural world. In this study, we propose a computational model to quantitatively derive motion speed across different spatiotemporal frequency bands in natural visual stimuli. This model was built and improved based on the well-known *motion energy* model and some of its extensions and variants.(Adelson and Bergen, 1985; Grzywacz and Yuille, 1990; Ogata and Sato, 1991; Etienne-Cummings et al., 1999; Shi and Luo, 2016) Such models employed multiple channels of spatial and temporal narrow-band filters to capture and synthesize the motion information (Bex and Dakin, 2002) (comprehensive reviews on the motion energy model are available (Burr and Thompson, 2011; Nishida, 2011)). Using an improved motion energy algorithm developed by us,(Shi and Luo, 2016) we were able to quantitatively verify the roles of different spatial frequency channels in visual motion speed perception in natural and stochastic (noise) image sequences by simulating different visual acuity levels such as normal vision (20/20), low vision (20/50 and 20/200, and complementary vision conditions containing only high frequency components.

## Methods

An overview of our methods for this work is shown in Figure 1. First, we generated motion sequences by translating different natural and stochastic images with known speed (Figure 2). Next, we filtered these sequences to simulate effects of different vision conditions: low-vision or loss of visual acuities at different levels using low pass filters with different cut-off frequencies and complementary vision conditions (hypothetical) using high pass filters (Figure 3). Spatiotemporally white noise was added to the sequences before the vision condition filtering, to simulate external (physical world) noise. Finally, we applied the biological motion perception model (Figures 1, 4 and 5) to estimate the speed of motion in these sequences (a variant of the one described in (Shi and Luo, 2016)). With the speed estimation results, we examined the relationship between spatial frequencies and motion perception accuracy, under different simulated vision conditions and at different speeds.

**Figure 1.**
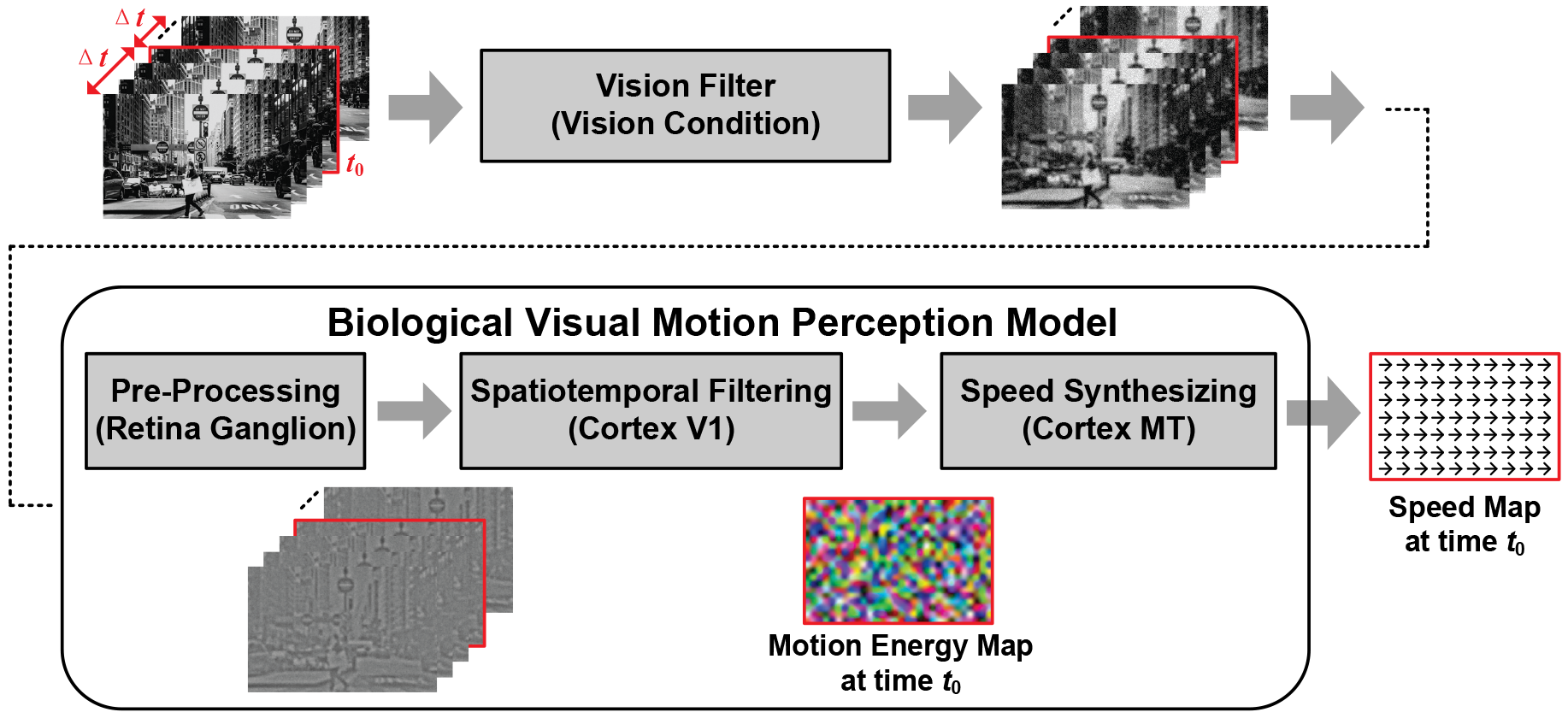
An overview of the motion energy model used for speed estimation in motion sequences. motion sequences were generated by translating images with different speeds. We filtered these sequences to simulate effects of different vision conditions and then we applied the biological motion perception model to these sequences. Motion estimation at time *t*_0_ requires the frames within a time window of 2*Δt* centered at *t*_0_, where 2*Δt* is the length of temporal filters used in the biological model. The preprocessing stage filtered out very high frequency noise and spatial DC components. Then the spatiotemporal filtering stage sampled multiple spatiotemporal frequency components to generate motion energy maps. Finally, the speed synthesizing stage used the motion energy maps to infer the real motion speed that was compared to the known value to obtain speed estimation error at different speed values as well as for different simulated vision conditions.

**Figure 2.**
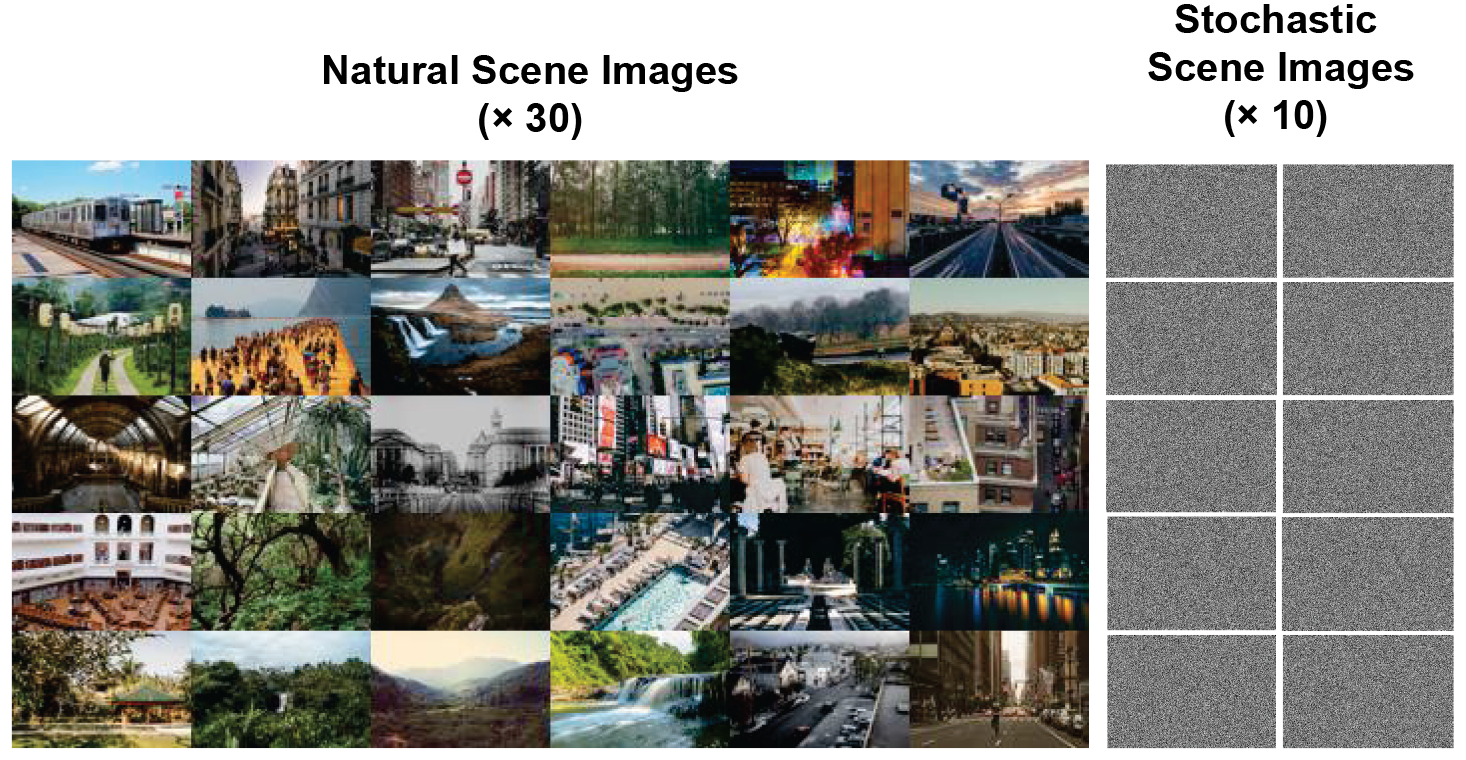
The high resolution daily life natural images (*n* = 30) downloaded from the internet and binary stochastic images (*n* = 10) were used for generating the motion sequences.

**Figure 3.**
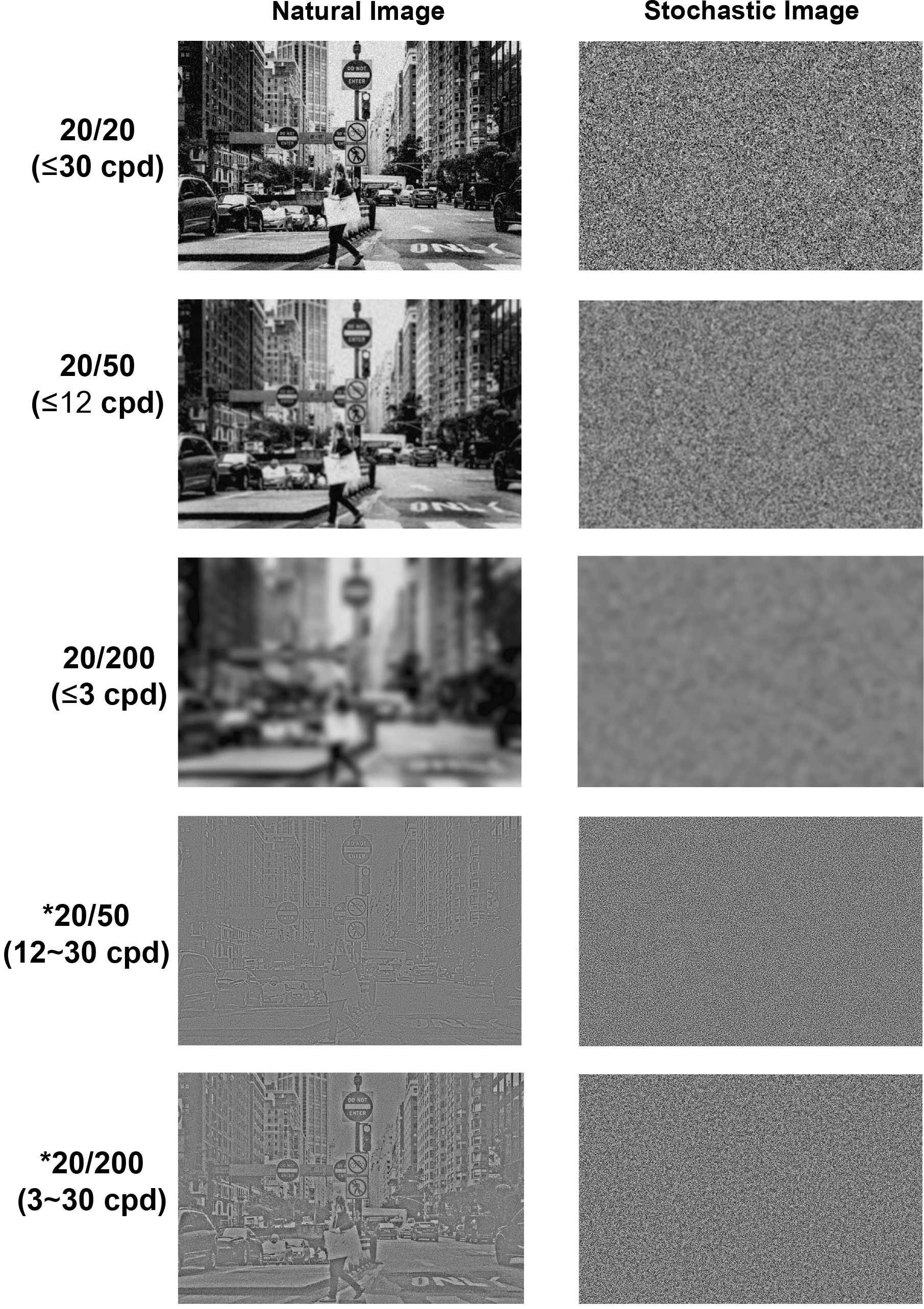
Simulated vision conditions. A frame from a natural sequence (left column) and a stochastic sequence (right column) filtered to simulate the 5 different vision conditions (each row). The visual acuity levels and the corresponding cutoff frequency is indicated for each row (* indicates complementary vision conditions). The low vision sequences only contained lower frequencies (blurring effect), whereas the complementary conditions, which do not exist in real world, contained only high frequency components (preserving edges).

### Generation of motion sequences

To generate motion sequences of natural scenes, we used Google image search engine with keywords like natural scenes, urban scenes, rural scenes, street blocks, buildings, railways, beaches and so on, and randomly picked 30 high resolution images out of the search results (Figure 2). These natural images were then down-sampled (with anti-aliasing processing) to 900 × 600 grayscale pixels to ensure that they contained sufficient high frequency components for simulating the 20/20 normal vision. To simulate the inherently continuous physical world, we set a high view-field resolution of 120 pixel/° on these images so that each such 900 × 600 image corresponded to a field-of-view of (900/120) × (600/120) = 7.5° × 5° in the physical world. To compare with low spatial frequency dominated natural images, we also generated 10 additional binary stochastic images of the same resolution in which each pixel was randomly assigned as either 0 (dark) or 1 (white) with equal probabilities. The spatial frequency spectrums of these stochastic images were nearly flat from low to high frequencies.

We generated motion sequences from the above 40 images by horizontally shifting them in a cyclic manner (the part shifted out on one side was to be shifted in on the other side). For each image, we generated 5 motion sequences with different speeds: 0.1, 1, 5, 15 and 30 °/s. The time duration of each sequence was set to 2Δ*t* = 0.2s, *exactly* covering the length of temporal filters employed in the biological motion perception model, as will be introduced later. Therefore, the estimated speed for a sequence was actually for the central time instant *t*_0_ = 0.1s (see Figure 1). Since the speed did not change over time and the image contents did not change despite the shift, the estimated motion at any frame was representative of the motion perception for the entire sequence. Thus, the need for using longer motion duration time for motion estimation was obviated.

### Simulation of vision conditions

By filtering the motion sequences in different manner, we simulated 5 vision conditions: visual acuity (VA) of 20/20 (normal vision), VA of 20/50 (moderate vision loss), VA of 20/200 (severe vision loss), a complementary VA of 20/50, and a complementary VA of 20/200. The underlying assumption in simulating these vision conditions was that low vision conditions correspond to loss of high frequency components in the scene.

On the other hand, the complementary vision conditions simulated a situation where only high frequency components were perceived. While this is not realistic, it helps us to investigate the causal relationship between frequency and motion speed perception. In terms of spatial frequency, a VA of 20/20 (normal vision) simulated the perceivable spatial frequencies up to 30 cycle/° (cpd) (Ref), while the 20/50 and 20/200 low vision conditions simulated perceivable frequencies no higher than 12 and 3 cpd, respectively. The normal and low vision conditions were realized by applying low-pass spatial filters (called vision filters in this paper) with cutoff frequencies of 30, 12 and 3 cpd, respectively, to every frame of the original motion sequences. The complementary vision conditions that simulated the perception of only high frequency components were realized by subtracting their corresponding low vision sequences from the normal vision sequence frame by frame. For example, the complementary VA of 20/50 simulated a condition where frequencies within between 12 and 30 cpd were perceptible. These imaginary complementary vision conditions that simulate perception of only high spatial frequencies do not exist in the real world. They were used as hypothetical references for studying the effects of low and high frequencies on motion perception.

Moreover, to simulate external noise in the physical world, we added spatiotemporally white Gaussian noises with a standard deviation of 10% of the dynamic range of image brightness to all sequences both before the vision condition filtering. The noise level amounts to 25.6 for the 256 gray-level images used in our simulation. Figure 3 shows the first frames of one natural motion sequence and one stochastic motion sequence after being filtered to simulate the 5 vision conditions, all with the simulated 10% external noise.

### Biological visual motion perception model

In this work, we use an improved variant of the original Adelson & Bergen’s motion energy model to explore the relationship between spatial frequency bands and visual motion perception [C. Shi, TCSVT 2018]. The improved model decomposed the estimation of 2D motion vector into independent estimations of two speed components in orthogonal (horizontal and vertical) directions, thereby significantly reducing the computational complexity. The principle of visual motion perception is illustrated in Figure 4, where we can estimate the speed *v*_*x*_ of horizontally moving sinewave grating from the unique spatiotemporal frequency couple (*f*_*x*_, *f*_*t*_) with *f*_*y*_ = 0. We call *V*_*x*_ as the characteristic speed of the spatiotemporal frequency couple (*f*_*x*_, *f*_*t*_). For a complicated motion scenario with multiple spatiotemporal frequency components, we sample some (*f*_*x*_, *f*_*t*_) components (all with *f*_*y*_ = 0), and use their characteristic speeds to estimate the real horizontal speed component. Similarly, we sample different (*f*_*y*_, *f*_*t*_) components (all with *f*_*x*_ = 0), and use their characteristic speeds to estimate the real vertical speed component. The horizontal and vertical speed components together form the motion speed vector.

**Figure 4.**
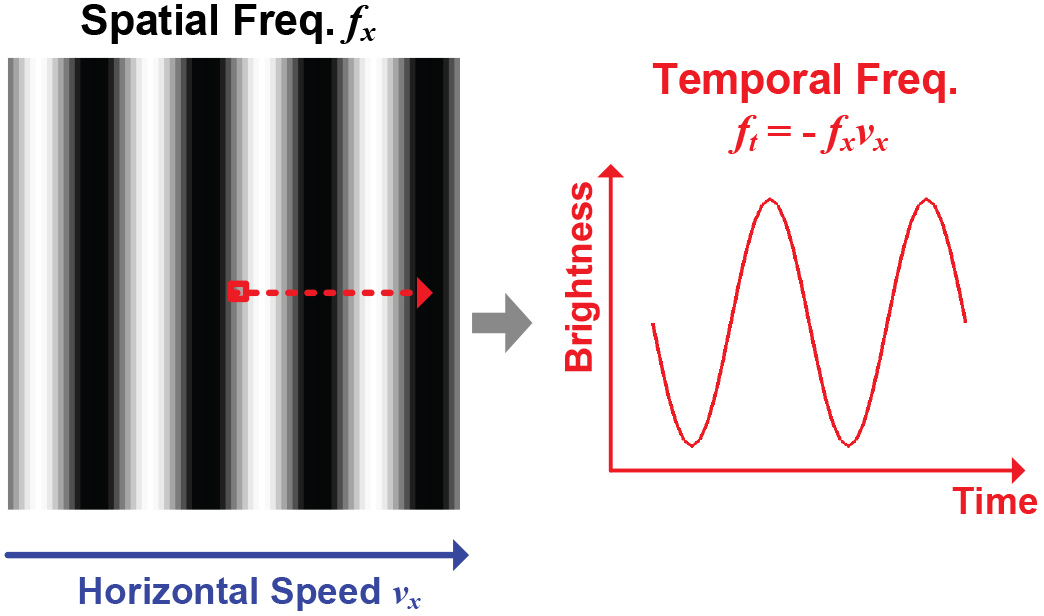
The relationship among spatial frequency *f*_*x*_, temporal frequency *f*_*t*_ and motion speed *v*_*x*_ at a an infinitesimally small location (shown as the red square on the sine-wave grating) during a short period of time. The horizontal motion of the sine-wave grating with a horizontal spatial frequency *f*_*x*_ (its vertical spatial frequency *f*_*y*_ is 0) invokes a temporal sine-wave of frequency *f*_*t*_ = −*v*_*x*_*f*_*x*_ at that location. So we could infer the horizontal speed *v*_*x*_ from the unique spatiotemporal frequency couple (*f*_*x*_, *f*_*t*_) of this simple stimulus where *f*_*y*_ is 0.

The biological motion perception model consists of three main stages that mimic the function of the primate visual pathway (see Figure 1):

1. Preprocessing stage. This stage removes high spatial frequency noise and spatial DC components in motion images, and only keeps textural features such as edges that are useful clues in motion perception. It is implemented by applying a wideband bandpass Difference-of-Gaussian (DoG) filter to every frame of the motion sequence. We set the pass band of the filter from 0.5 to 36 cpd in the simulation.
2. Spatiotemporal filtering stage. This stage extracts spatiotemporal frequency components called motion energies from the motion sequence (Figure 5). The preprocessed sequence is first spatially filtered by convolving with a group of spatial Gabor filters of different horizontal frequencies *f*_*x*_ (their vertical frequencies *f*_*y*_ are all 0). Then each resulting image is temporally filtered by convolving with another group of temporal Gabor filters of different temporal frequencies, pixel by pixel along the time dimension. Since the motion sequences in our simulation had the same length as the temporal filters, each temporally filtered result would be only one image corresponding to the central time *t*_0_. The values along the spatial locations for this resultant image are just the spatiotemporal frequency components, or motion energies. The motion energy for a given spatial image location across all the motion energy maps corresponding to different speeds can be concatenated as a motion vector, after which the local horizontal speed component could be inferred. For vertical speed component, this stage uses spatial Gabor filters of different vertical frequencies *f*_*y*_ (their horizontal frequencies *f*_*x*_ are all 0), which can be obtained by rotating the spatial Gabor filters in Figure 5 by 90°. In our simulation, the sampled spatial frequencies for spatial Gabor filters are 0.6m cpd, *m* = 1, 2, …, 50. And the sampled temporal frequencies for temporal Gabor filters are 5m cycle/s (Hz), *m* = −10, −9, …, −1, 0, 1, …, 9, 10. Here the maximum absolute value of sampled temporal frequencies is 50 Hz, which comes from the limit of human visual system. With the relationship *f*_*t*_ = −*v*_*x*_*f*_*x*_ as mentioned in Figure 4, we need to sample both positive and negative temporal frequencies to distinguish speed directions (rightward/leftward or upward/downward for the horizontal or vertical speed component, respectively).
3. Speed synthesizing stage. Figure 6 shows how motion energies are used to infer the motion speed in this stage. The motion energy of a spatiotemporal couple (*f*_*x*_, *f*_*t*_) represents a Gaussian distribution of speed likelihood with its mean at the characteristic speed (− *f*_*t*_ / *f*_*x*_) (Grzywacz, N. M. 1990) given by:

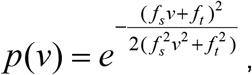

where *f*_*s*_ denotes either horizontal or vertical spatial frequency (*f*_*x*_ or *f*_*y*_), depending on whether the horizontal or vertical speed component (*v*_*x*_ or *v*_*y*_) is being estimated. To estimate the horizontal or vertical speed component at an image location, these speed likelihood distributions are weighted by the values of their corresponding motion energies of sampled spatiotemporal frequencies at that location, and then accumulated. The motion speed at that location is estimated as the one with the highest accumulated likelihood. Since we simulated 5 motion speeds from 0.1 to 30 °/s, we calculated likelihoods on discrete speed samples within a range from −40 to 40 °/s, with a step of 0.01 °/s from −0.2 to 0.2 °/s, and a step of 0.1 °/s elsewhere. Such range and small steps were designed to reduce the impact of quantization on motion perception accuracy as much as possible.

**Figure 5.**
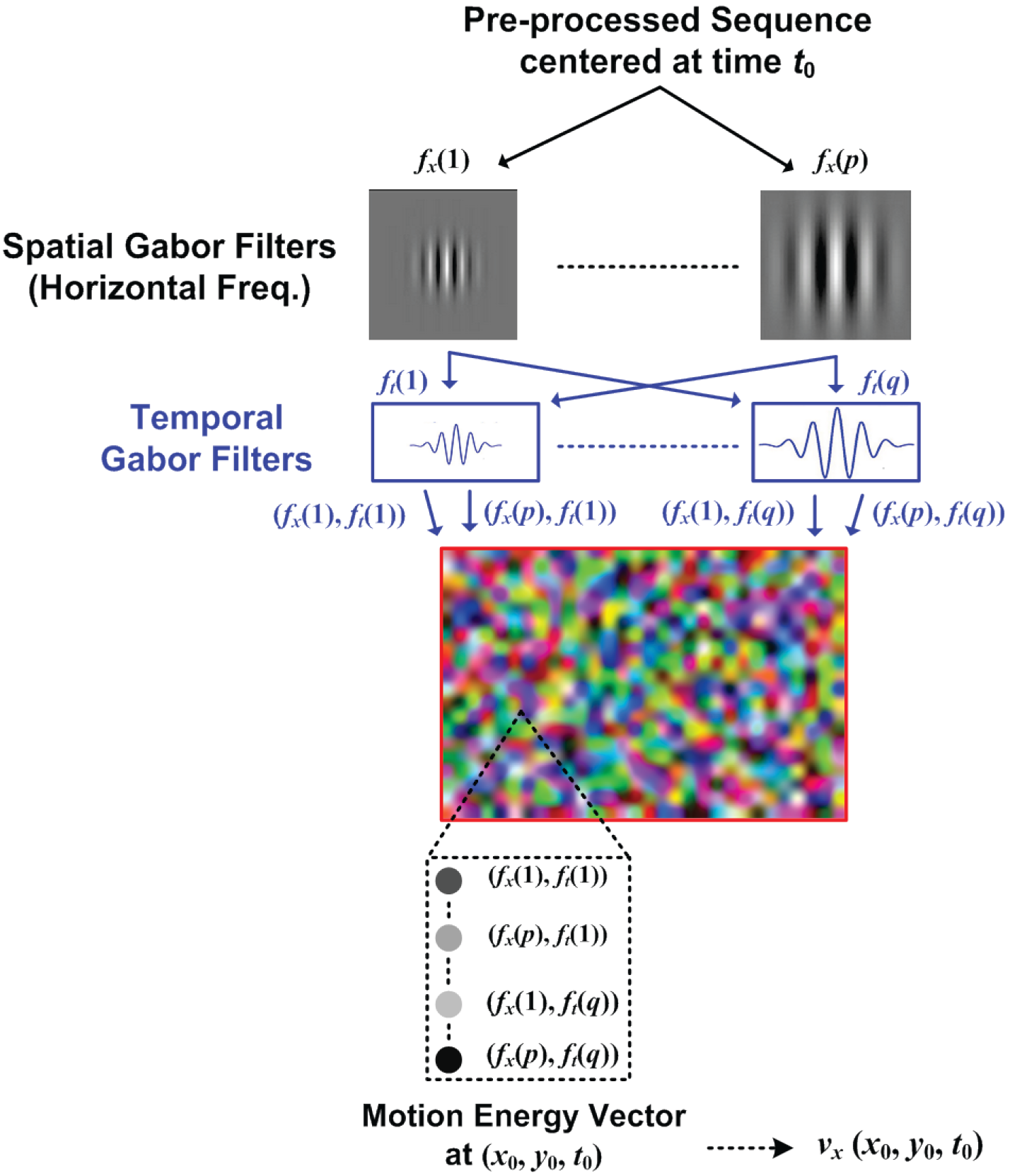
The spatiotemporal filtering stage in our biological motion perception model. The preprocessed sequence is first convolved with a group of spatial Gabor filters of different horizontal frequencies *f*_*x*_ (their vertical frequencies *f*_*y*_ are all 0). Then each resulting image is convolved with another group of temporal Gabor filters. Thus the spatiotemporal frequency components of the motion sequence are extracted as the motion energy maps. The horizontal speed component at a spatial image location could be inferred from the motion energy values at that location. For vertical speed component, this stage uses spatial Gabor filters of different vertical frequencies *f*_*y*_ (their horizontal frequencies *f*_*x*_ are all 0), which can be obtained by rotating the spatial Gabor filters by 90°.

**Figure 6.**
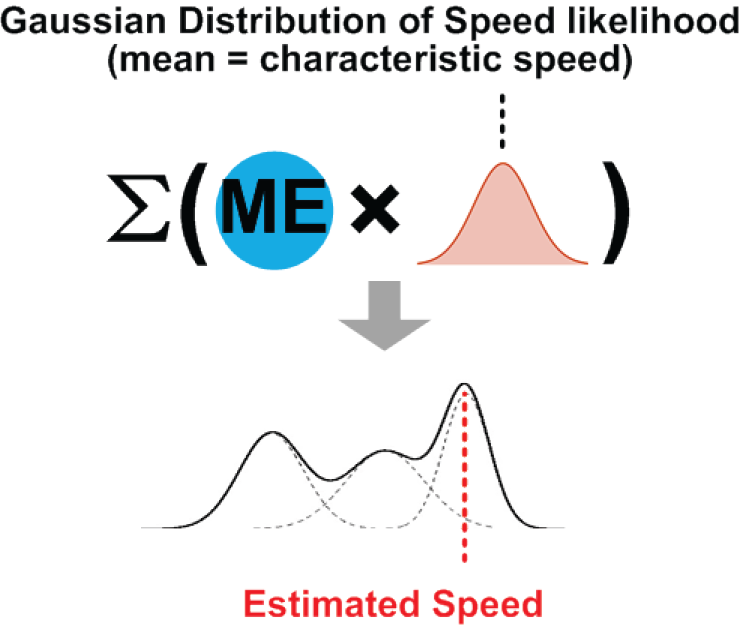
The speed synthesizing stage in the biological motion perception model. The motion energy of a spatiotemporal couple (*f*_*x*_, *f*_*t*_) represents a Gaussian distribution of speed likelihood with its mean at the characteristic speed (− *f*_*t*_ / *f*_*x*_). For most natural image sequences, there are many spatiotemporal couples present at any given image location. To estimate the horizontal or vertical speed component at a location, these speed likelihood distributions are accumulated after weighting by the values of their corresponding motion energies of sampled spatiotemporal frequencies. The motion speed at that location is estimated as the one with the highest accumulated likelihood (winner take all strategy).

The estimation of horizontal and vertical speed components is carried out separately and independently in this biological model, except using spatial Gabor filters in different orientations as mentioned before. Moreover, all spatial locations of simulated motion sequences saw the same speed under the horizontal shifting. To accelerate simulation, we selected probing locations with an interval of 20 pixels (1/6 ° in view field) along each image dimension (excluding locations near the boundary lying within the radius of spatial Gabor filters), and used the average estimated speed on those locations as the perceived speed for the entire sequence.

### Evaluation and Statistical Analysis

We present the results of our simulation in terms of motion energy distributions for different vision conditions for natural and stochastic motion sequences. For a given speed and vision condition, the distributions were obtained by accumulating the motion energy for each sampled spatial frequency (with the temporal frequencies integrated out) for all the probing spatial image locations and then averaged over all the 30 natural sequences. We then present the speed estimation results in terms of relative error in motion perception compared to the ground truth. We performed repeated measures anova to determine the within-subject effects of different speeds and vision conditions on speed estimation error for a given type of image sequence, and between subject effects of sequence type (each image sequence can be considered to be a subject). The relative speed estimation errors were inverted for natural sequences to ensure normality of the data; the data for stochastic sequences were normal (normality tested using Shapiro-Wilks test). Non parametric testing method was used for comparing the speed estimation error for natural and stochastic sequences (2 sample Kolmogorov-Smirnov test). Statistical analysis was performed using IBM-SPSS.

## Results

The raw motion energy distributions over spatial frequencies in natural sequences for different speeds in normal vision condition are highly skewed toward low spatial frequencies (Figure 7a). The normalized cumulative motion energy distributions level-off in the low frequency range, further clarifying the concentration of the motion energy near the low spatial frequencies (Figure 7b). For instance, about 90% of the total motion energy is concentrated below 12 cpd frequencies for 0.1°/s speed curve and this amount increases for other speed curves, going up to 99.9% for 30°/s speed. Due to the higher concentration of motion energy at lower frequencies in natural sequences, there is also a relatively large overlap between the motion energy distributions for normal vision and the low vision conditions (85% for VA 20/50 and 45% for 20/200), but relatively smaller overlap for the complementary vision conditions (38% for VA *20/200 and 3% for *20/50). Larger amount of overlap indicates more similarity in the motion energy distributions. Since the cutoff frequencies for vision condition simulating filters are 3 and 12 cpd (the low vision conditions are low pass cases, whereas the complementary conditions are high pass cases around these cutoff frequencies), larger overlap of normal vision and low vision conditions is the expected outcome.

**Figure 7.**
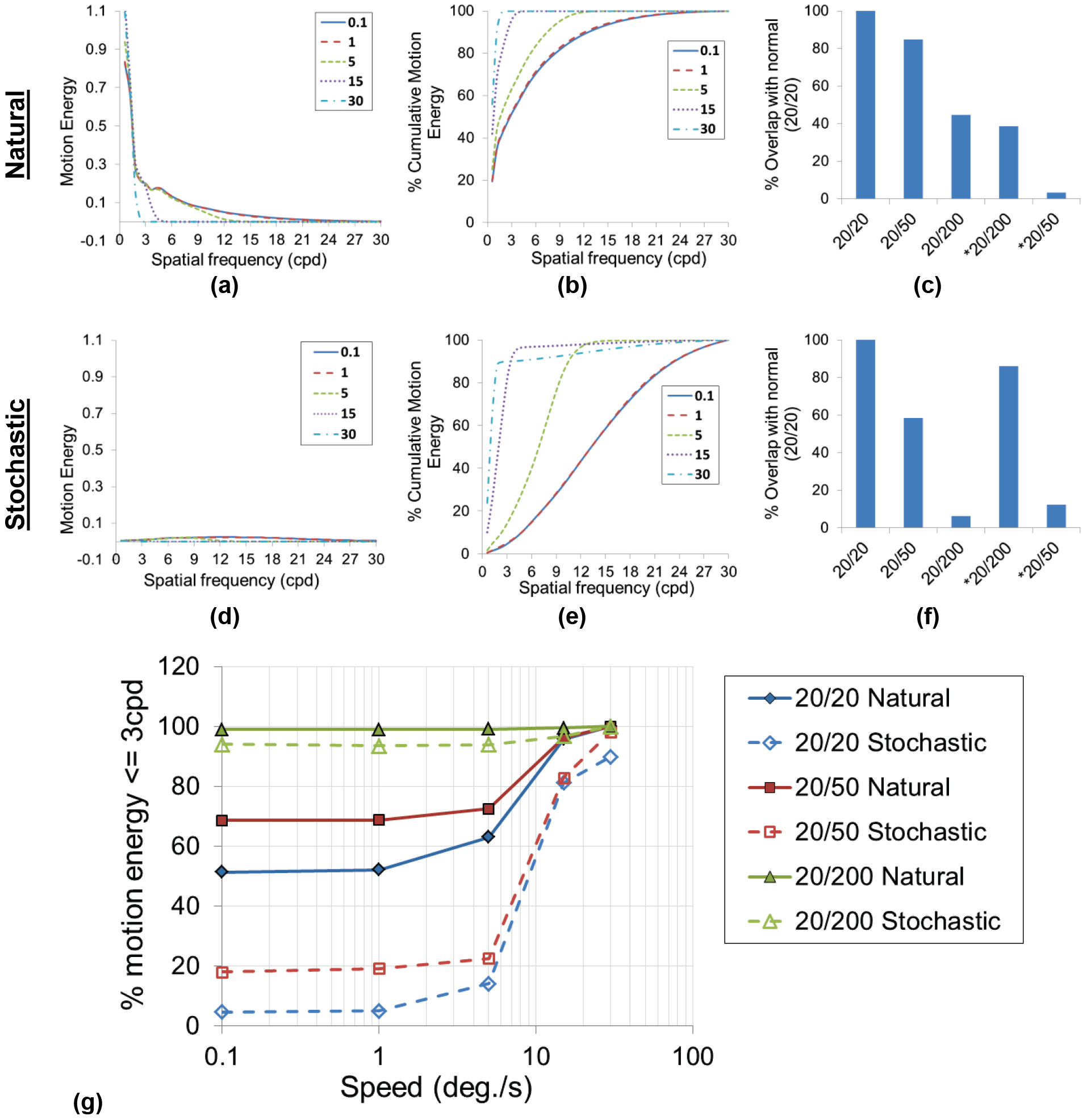
Motion energy distribution over spatial frequency in normal vision condition for different speeds in natural (a) and stochastic (d) sequences. Normalized cumulative motion energy curves in normal vision conditions for different speeds in natural (b) and stochastic (e) sequences. Percent overlap of motion energy distributions for various vision conditions (low-vision and complementary vision conditions) with normal vision (20/20) in natural (c) and stochastic (f) sequences. A larger overlap means higher degree of similarity between the distributions. Median speed (5°/s) curve was used for computing the overlap. Effect of speed on the motion energy distributions for normal and low vision conditions in natural and stochastic sequences (g). The fraction of motion energy below 3cpd frequency indicates its concentration in low frequency region, which increases with increasing speed.

Compared to natural sequences, the motion energy distributions for stochastic images in normal vision condition are no longer concentrated near low frequency bands and instead appear flat across the entire frequency spectrum (Figure 7d). The motion energies are also of significantly smaller magnitude (reduced by about 89%). The relatively flat motion energy distribution in stochastic sequences is also evidenced by the relatively slow rising 0.1°/s speed curve in Figure 7e (40% motion energy concentrated below 12cpd compared to about 90% for natural sequences for the same frequency band). Furthermore, contrary to the natural sequences, there is a relatively larger overlap between normal vision and complementary vision conditions (86% for *20/200 and 12% for *20/50) than the low vision condition (58% for 20/50, 6% for 20/200) (Figure 7f). Again, this is expected since there is relatively small amount of motion energy present in the low frequency band in the stochastic sequences to begin with.

There is a discernable effect of speed on the motion energy distributions, as higher speeds lead to a higher concentration of motion energy in the lower frequency region in both natural and stochastic sequences (Figure 7g). Since 3 cpd was the lowest cutoff frequency for simulation of low vision conditions, the fraction of motion energy at or below 3 cpd was used as a way to quantify the effect of speed on motion energy distributions. Predictably for the 20/200 vision condition with 3 cpd cutoff frequency, the motion energy fraction at 3cpd is already at 99% at 0.1°\s. In natural sequences for normal vision and 20/50 condition, the fraction of motion energy ≤ 3cpd at 0.1°/s speed is at 51% and 68%, respectively. As the speed increases, this amount increases close to about 100% at 30°/s. The same effect is also seen in stochastic sequences. However, the motion energy fraction ≤ 3 cpd for 0.1°/s is a lot lower in stochastic sequences compared to natural sequences (5% and 18% for 20/20 and 20/50 condition for stochastic, whereas 51 and 68% for the same in natural sequences), before increasing to close to 100% at 30°/s. There is also a noticeable interaction of vision conditions and speed on the motion energy distribution within natural and stochastic sequences: lower visual acuity leads to less steeper increase in motion energy fraction ≤ 3 cpd for higher speeds. This is again expected, since more motion energy is concentrated in lower frequency regions for low vision conditions to begin with.

Motion perception in low vision condition was not significantly affected in natural sequences as the speed estimation error did not change significantly in normal vision and the 2 low vision conditions (*F* = 2.745, *p* = 0.099) (Figure 8a). There was a significant effect of speed as the error increased with speed in all vision conditions (*F* = 34.92, *p* < 0.001). The interaction between speed and vision conditions was significant (*F* = 4.39, *p* = 0.034). In contrast to natural sequences, speed estimation error was significantly larger in 20/20 case compared to other low vision conditions in stochastic sequences (*F* = 849.2, *p* < 0.001) (Figure 8b). The error also significantly increased with speed in stochastic sequences (*F* = 1029.34, *p* < 0.001). The interaction of speed with vision conditions was significant (*F* = 21.39, *p* < 0.001), as the error in 20/20 condition was significantly higher than 20/50 and 20/200 vision conditions at higher speeds. Speed estimation was impaired in complementary vision conditions as the error was significantly larger in complementary conditions compared to the normal vision or low vision conditions (*F* = 26.559, *p* < 0.001) (Figure 8c). The error increased significantly with speed in the complementary vision conditions (*F* = 106.13, *p* < 0.001) and the interaction between speed and vision conditions was significant (*F* = 21.39, *p* < 0.001) as the error between complementary and low vision conditions was larger at higher speeds. When comparing the two kinds of sequences in normal and low vision conditions, the speeds estimation error was significantly different in natural and stochastic sequences (*F* = 9.23, *p* = 0.005) (Figure 8d). There was a significant differences in error distribution (pooled across all speeds) in all three vision conditions between natural and stochastic sequences (20/20: *Z* = 2.32, *p* < 0.001; 20/50: *Z* = 1.94, *p* = 0.001; 20/200: *Z* = 3.01, *p* < 0.001), even as the medians did not differ. This was due to highly skewed nature of the error distribution, which necessitated use of non-parametric statistical approach in this particular case.

**Figure 8.**
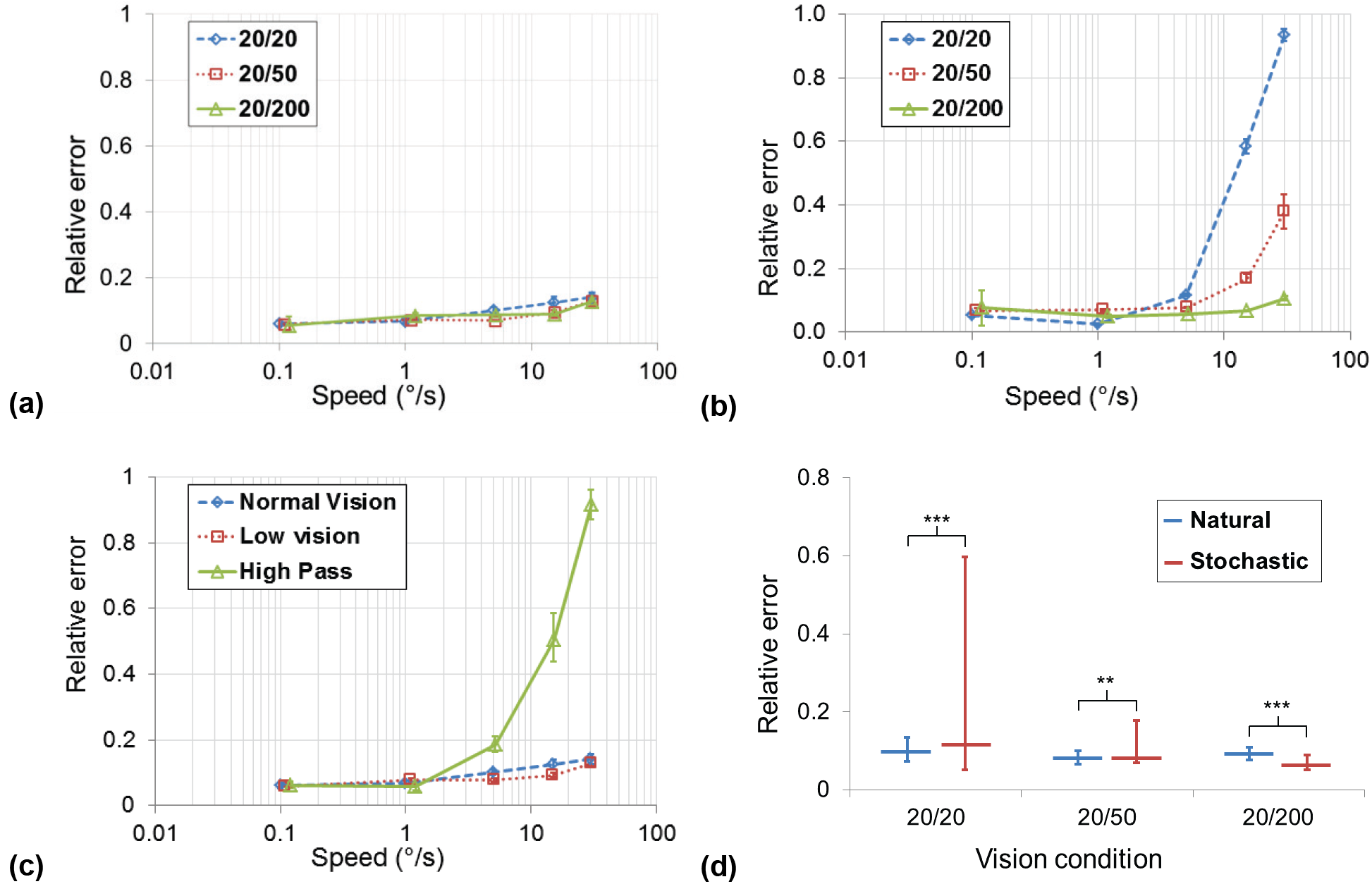
Speed estimation error results. Comparison of relative error (ratio of absolute error and ground truth speed) in normal vision (20/20) and low vision conditions (20/50 and 20/200) over the tested speed range is shown for natural (a) and stochastic (b) sequences. Comparison of relative speed estimation error between normal vision, low vision, and complementary vision conditions (high pass) in natural sequences are shown in (c). Low vision condition in this case is the mean of 20/50 and 20/200 conditions, whereas the high pass condition is the mean of *20/50 and *20/200 conditions. In plots (a) through (c), error bars denote standard error of mean. Also in these plots please note the logarithmic scale for the horizontal axis. Comparison of the relative error for natural and stochastic sequences is shown in (d). The pooled relative error distributions across all speeds for 20/20, 20/50, and 20/200 conditions are shown, with horizontal lines representing the median and the error bars showing the 25^th^ and 75^th^ percentile (***: *p* < 0.001, **: 0.01 < *p* ≤ 0.001).

## Discussion

We have presented an analytical model based on motion energy to examine the relationship between motion perception and spatial frequency. Direct speed estimation is one of the main features of our models, providing a more objective way of determining motion perception sensitivity compared to motion perception based psychophysical thresholds, as done in previous human subject studies. By simulating different vision conditions, we explored the causal relationships between motion perception accuracy and spatial frequency in natural images. Overall, our results show the dominant role played by low spatial frequency components in motion perception that largely agree with a wide variety of previous research in motion psychophysics and primate neurobiology.

### Spatial frequency and motion perception

In this work we simulated three broad categories of vision conditions: normal vision (20/20), low vision via low pass filtering (20/50 and 20/200), and complementary high pass vision conditions (*20/50 and *20/200). The speed estimation results (Figure 8) showed two things: i) as long as there are low spatial frequencies in the images, speed estimation is equally accurate with and without high spatial frequencies, and ii) speed estimation suffers with large errors when there are only high spatial frequencies (see high pass filtered complementary vision conditions). Together, these suggest that low frequency components are key for motion perception. Specifically, perception of motion at higher speeds requires lower spatial frequencies, as seen in the biasing of motion energy curves toward low frequency bands for higher speeds in Figure 7. This is consistent with the findings previously reported in different forms and contexts in a number of studies over a span of about three decades.

In many psychophysical studies, increasing speeds led to a shift or biasing of the response toward low frequencies. Vernier acuity thresholds were shown to reduce with increasing spatial frequencies of the stimulus presented at different speeds.(Chung et al., 1996; Levi, 1996; Mechler and Victor, 2000) For a given spatial frequency value, the Vernier thresholds were shown to have a characteristics relationship with velocities: thresholds remained constant for a range of velocities before decreasing linearly at a velocity value known as the “knee point” of the curve. This “knee point” shifted toward lower spatial frequencies with increasing velocities, indicating the inability to perceive high frequency components at higher speeds.

Global motion coherence thresholds were shown to follow a U shaped curve as a function of spatial frequency, and the frequency of highest sensitivity (valley of the U shaped curve) reduced with increase in speed.(Hess and Aaen-Stockdale, 2008) Similar effects of spatial frequency on Weber fraction thresholds were shown for velocity discrimination tasks, where increasing speeds led to a progressive decline in the spatial frequency for highest motion sensitivity.(McKee et al., 1986; Chen et al., 1998)

No effect of high spatial frequencies was found in motion detection, as detection rates did not differ for blurred stimuli (low spatial frequency) vs. spatially bandpass stimuli (low and high frequency content).(Wichmann and Henning, 1998) In a texture segregation task using random-dot kinematogram, the maximum distance of movement for which the texture could be perceived was smaller for low frequency stimuli compared to high frequency stimuli.(Gilden et al., 1990) Direction of apparent motion was shown to be determined by low frequencies in a task where a stationary square appeared to move toward a low pass flanking stimuli.(Ramachandran et al., 1983)

Some studies presented raw speed thresholds over a range of spatial frequencies. (Yang and Stevenson, 1997; Shioiri et al., 2002) Particularly, Yang & Stevenson show the lowest speed thresholds are obtained for spatial frequencies close to 2 cpd, with a slight increase in the thresholds for larger frequencies (data is limited at 12 cpd). However, they also show an increase in speed thresholds for very low frequencies, indicating the limitation of motion perception at very low spatial frequencies. While our results are trending in a similar manner (increase in relative speed estimation error for 0.1 deg/s speeds in Figure 6), limitations of simulation setup precludes us from evaluation at lower spatial frequencies. This point is elaborated in detail below.

Despite the underlying methodological differences, collectively, these previous studies show the relatively reduced role of higher frequencies and an increased role of low frequencies in motion perception with higher motion speeds. It should be noted that many of the above studies reported motion perception by human observers in terms of psychometric thresholds or detection rates.(Ramachandran et al., 1983; Gilden et al., 1990; Wichmann and Henning, 1998; Hess and Aaen-Stockdale, 2008) Our analytical model, on the other hand, allows direct estimation of perceived speeds and thus our evaluation consisted of speed estimation errors for a large range of motion speeds. Direct speed estimation for slow and fast motion is more relevant for real world tasks such as driving or walking, where we need to see not only what is moving but also estimate the speeds in order to avoid collisions or plan an appropriate path for navigation.

While we have shown that our simulation results of the motion perception model are consistent with the previous work, the question remains: why high frequency information is irrelevant in motion perception? A straightforward explanation is embedded in the relationship of spatial frequency, temporal frequency, and speed (as shown in Figure 1). For a given temporal frequency, the speed is inversely proportional to the spatial frequency. At high speeds and high spatial frequencies, the limit of temporal frequency is reached. (Nakayama, 1990) This was explained in terms of spatiotemporal motion energy units.(Yang and Stevenson, 1997) Considering the spatiotemporal motion detector field with spatial frequency on x axis and temporal frequency on the y axis, the slope represents the speed of detected motion. Now, for any given spatial and temporal frequency pair, there is an upper bound on the slope i.e., the speed that can be perceived, and this upper bound reduces (largest speed that can be perceived becomes smaller) when spatial frequency is reduced while keeping temporal frequency constant. Thus, for mid to higher speed range (above 1 °/s) the role of higher spatial frequencies will be progressively reduced and their elimination will not lead to motion perception deficits for larger motion speeds.

### Motion perception and low vision

We hypothesized that loss of visual acuity may not result in significant motion perception deficits based on the following rationale: i) loss of visual acuity generally corresponds to the inability to perceive high frequency components and ii) high frequency components do not play a key role in motion perception compared to low frequency components (as discussed above). Evaluation of our motion perception model support this hypothesis as motion energy distribution (Figure 7) and speed estimation errors (Figure 8) for low vision conditions are similar to normal vision condition.

From visual pathway perspective, motion perception and object vision processing are considered to occur along separate processing pathways in the brain, with motion being perceived via dorsal stream and object recognition being processed via ventral stream.(Goodale et al., 1991) While the two pathways are interconnected, deficits in one pathway may not necessarily affect the other pathway, during early development (childhood) or in aging.(Owsley, 2011; Chakraborty et al., 2015; Hadad et al., 2015)

Psychophysical studies have explored the relationship of spatial frequency and low vision motion perception using simple visual tasks. Pan & Bingham showed that motion information facilitated detection of events in highly blurred videos.(Pan and Bingham, 2013) Lappin et al. showed that motion detection thresholds in low vision subjects were similar to normally sighted subjects for speeds > 2 degrees/second for foveal viewing.(Lappin et al., 2009) Similarly, Tadin et al. showed that there were no adverse effects of visual acuity loss on motion perception for low spatial frequency stimuli.(Tadin et al., 2012) Studies involving real-world motion perception tasks have also found similar trends regarding the independence of motion perception and visual acuity loss. For instance, motion detection sensitivity was the only independent predictor of performance in driving related tasks in the elderly and in AMD subjects, irrespective of their VA and CS scores.(Wood et al., 2014; Wood et al., 2018) Thus, central vision loss (loss of visual acuity) may not result in significant degradation in driving performance, which is correlated with motion sensitivity that is independent of visual acuity. Saunders et al. found no significant differences in subjects’ description of then events when viewing blurred videos simulating VA loss compared to when viewing videos without blur.(Saunders et al., 2014)

Our results based on motion energy distribution in different vision conditions over various spatial frequency bands reinforce the finding of the human subject studies and provide an analytical framework to explain why loss of visual acuity may not significantly affect motion perception tasks. However, this does not imply that people with visual impairments will be as perfect as normally sighted people in all motion perception tasks because there are caveats to the above mentioned studies and observations. While motion perception was not affected by visual acuity loss for foveal viewing, impairment may occur in both central and peripheral vision at the same in some visually impaired people.(Lappin et al., 2009; Tadin et al., 2012) Similarly, while VA was not a factor predicting driving performance, Wood el al. did find a significant degradation in performance in AMD subject group compared to age-matched normally sighted subjects. Also, even though low vision means impaired ability to see high frequency components, in reality, perception of a wider band of frequencies may be hampered.(Chung and Legge, 2016) Because low vision can manifest in various different forms, it is challenging to characterize it with a single metric. We believe that, in order to understand visually impaired people’s performance for real world tasks, both static (object) vision such as VA and CS and motion perception need to be taken into consideration.

### Natural images vs. artificial stimuli

One of the key differences in our analysis compared to the vast majority of previous motion perception studies was the use of natural motion sequences. Importance of using natural stimuli as opposed to artificial stimuli in understanding vision has been noted recently. (Kayser et al., 2004; Felsen and Dan, 2005; Einhauser and Konig, 2010) Particularly in the context of motion perception, the choice of stimuli can affect the outcomes. Main reason for this is that the spatial frequency distribution in the natural images tends to be different than artificial stimuli, which can be critical when studying motion perception related to real world tasks. Spectrums of natural images are typically broadband, but dominated by low frequency components,(van der Schaff and van Hateren, 1996) whereas artificial stimuli used in previous studies are often narrow band, e.g. Gabor patches, or flat band, e.g. random dots. Although they are related, we should point out that motion energy is not equivalent to the power spectrum of an image. While spectral signatures were found to be different in different categories of natural images such as nature landscape and urban scenes,(Torralba and Oliva, 2003) there was little difference in motion energy distribution in these images in our experiments.

Using stochastic sequences, we demonstrate how the presence of a relatively flat spatial frequency distribution that was very much unlike the natural images can affect motion perception. An interesting finding is that normal vision produced larger errors at high speeds (15, 30 °/sec) than the low vision conditions (20/50 and 20/200) (Figure 8c). This can be explained by the low signal-to-noise ratio (SNR) of stochastic sequences.

As we showed above, speed estimation for fast motion is highly reliant on the motion energy extracted from low spatial frequency bands. For stochastic sequences with added noise, the SNR in the low frequency band was much lower compared to the natural sequences, which led to higher speed estimation errors in normal vision condition. When the sequences were low pass filtered to simulate 20/50 and 20/200 vision conditions, the SNR improved and allowed more accurate speed estimation compared to NV condition. For both stochastic and natural scenes, the SNR at low frequency band is even lower in the *20/50 and *20/200 conditions, therefore resulting in large errors for high speeds (Figure 8). This finding is in line with Van Droon’s study, which found a high SNR is required for accurate speed estimation in the case of fast visual motion.(Van Doorn and Koenderink, 1982) Further psychophysics studies are needed to confirm this simulation results in human observers.

### Limitations

The simulations of low-vision conditions used for evaluation in this work have some limitations. First, extremely low visual acuities (below 20/200) were not simulated as they required a very high spatiotemporal resolutions and large filter sizes in order to satisfy the Nyquist rate. At very low spatial frequencies, we expect motion sensitivity be impaired for slow speeds, (Yang and Stevenson, 1997) which means that people with very low visual acuities will not be able to perceive slow motion speeds. Due to the limits of the Nyquist sampling rate, we were not able to simulate ultra-low visual acuity cases to try and reproduce this result (our spatial frequencies were limited to 0.6 cpd, whereas Yang & Stevenson went as low as 0.25 cpd).

Another limitation of the simulation framework relates to the assumption that low vision conditions correspond to inability to perceive higher frequencies. As described earlier, in real world, low vision corresponds to a variety of conditions where the perception of a very wide range of spatial frequencies can be impaired in a non-uniform manner.(Chung and Legge, 2016) Thus, our simulation of low vision condition is only approximate representation of real-world cases.

Another limitation in the motion perception model could be caused due to separable implementation of horizontal and vertical speed estimation components. While such an implementation makes it an efficient algorithm, in real-world speed perception is performed in scenes where motion is present in multiple directions. Presence of multiple simultaneous motion components may degrade speed estimation performance.(Gal et al., 2009; St.Clair et al., 2010) Tweaking the motion perception model for speed estimation in presence of orthogonal distractors is future work.

## References

Adelson EH, Bergen JR (1985) Spatiotemporal energy models for the perception of motion. Journal of the Optical Society of America - A 2:284–299.

Bex PJ, Dakin SC (2002) Comparison of the spatial-frequency selectivity of local and global motion detectors. Journal of the Optical Society of America A-Optics Image Science and Vision 19:670–677.

Borst A, Euler T (2011) Seeing things in motion: models, circuits, and mechanisms. Neuron 71:974–994.

Burr D, Thompson P (2011) Motion psychophysics: 1985-2010. Vision Research 51:1431–1456.

Chakraborty A, Anstice NS, Jacobs RJ, Paudel N, LaGasse LL, Lester BM, Wouldes TA, Harding JE, Thompson B (2015) Global motion perception is independent from contrast sensitivity for coherent motion direction discrimination and visual acuity in 4.5-yearold children. Vision Research 115:83–91.

Chen Y, Bedell HE, Frishman LJ (1998) The precision of velocity discrimination across spatial frequency. Perception & Psychophysics 60:1329–1336.

Chung S, Legge G (2016) Comparing the shape of contrast sensitivity functions for normal and low vision. Investigative Ophthalmology & Visual Science 57:198–207.

Chung STL, Levi DM, Bedell HE (1996) Vernier in Motion: What Accounts for Threshold Elevation? Vision Research 36:2395–2410.

Einhauser W, Konig P (2010) Getting real—sensory processing of natural stimuli. Current Opinion in Neurobiology 20:389–395.

Etienne-Cummings R, Van der Spiegel J, Mueller P (1999) Hardware implementation of a visual-motion pixel using oriented spatiotemporal neural filters. IEEE Transactions on Circuits and Systems II: Analog and Digital Signal Processing 46:1121–1136.

Felsen G, Dan Y (2005) A natural approach to studying vision. Nature Neuroscience 8:1643–1646.

Gal V, Kozak LR, Kobor I, Banko EM, Serences JT, Vidnyanszky Z (2009) Learning to filter out visual distractors. European Journal of Neuroscience 29:1723–1731.

Gilden DL, Bertenthal BI, Othman S (1990) Image Statistics and the Perception of Apparent Motion. Journal of Experimental Psychology 16:693–705.

Goodale M, Milner A, Jakobson L, Carey D (1991) A neurological dissociation between perceiving objects and grasping them. Nature 349:154–156.

Grzywacz NM, Yuille A (1990) A model for the estimate of local image velocity by cells in the visual cortex. Proceedings of the Royal Society of London B: Biological Sciences 239:129–161.

Hadad B, Schwartz S, Maurer D, Tl L (2015) Motion perceptions review of developmental changes and the role of early visual experience. Frontiers in Integrative Neuroscience 9.

Hess RF, Aaen-Stockdale C (2008) Global motion processing: The effect of spatial scale and eccentricity. Journal of Vision 8:1–11.

Kayser C, Kording KP, Konig P (2004) Processing of complex stimuli and natural scenes in the visual cortex. Current Opinion in Neurobiology 14:468–473.

Lappin JS, Tadin D, Nyquist JB, Corn AL (2009) Spatial and temporal limits of motion perception across variations in speed, eccentricity, and low vision. Journal of Vision 9:1–14.

Levi DM (1996) Pattern perception at high velocities. Current Biology 6:1020–1024.

Liu J, Newsome WT (2002) Functional Organization of Speed Tuned Neurons in Visual Area MT. Journal of Neurophysiology 89:246–256.

McKee S, Silverman G, Nakayama K (1986) Precise velocity discrimination despite random variations in temporal frequency and contrast. Vision Research 26:609–619.

Mechler F, Victor JD (2000) Comparison of thresholds for high-speed drifting vernier and a matched temporal phase-discrimination task. Vision Research 40:1839–1855.

Nakayama K (1990) Properties of early motion processing: Implications for the sensing of ego motion. In: The Perception and Control of Self Motion (Warren R, A.H. Wertheim, eds), pp 69–80. Hillsdale NJ: Lawrence Erlbaum.

Newsome WT, Pare EB (1988) A selective impairment of motion processing following lesions of the middle temporal visual area (MT). The Journal of Neuroscience 8:2201–2211.

Nishida S (2011) Advancement of motion psychophysics: Review 2001-2010. Journal of Vision 11:11.

Ogata M, Sato T (1991) Motion perception model with interaction between spatial frequency channels. Systems and computers in Japan 22:30–39.

Orchard G, Etienne-Cummings R (2014) Bio-inspired Visual Motion Estimation. Proceedings of the IEEE 102:1520–1536.

Owsley C (2011) Aging and vision. Vision Research 51:1610–1622.

Pan JS, Bingham GP (2013) With an Eye to Low Vision: Optic Flow Enables Perception Despite Image Blur. Optometry and Vision Science 90:1119–1127.

Perrone JA, Thiele A (2001) Speed skills: measuring the visual speed analyzing properties of primate MT neurons. Nature Neuroscience 4:526–532.

Priebe NJ, Lisberger SG, Movshon JA (2006) Tuning for Spatiotemporal Frequency and Speed in Directionally Selective Neurons of Macaque Striate Cortex. The Journal of Neuroscience 26:2941–2950.

Ramachandran VS, Ginsburg AP, Anstis SM (1983) Low Spatial-Frequencies Dominate Apparent Motion. Perception 12:457–461.

Saunders DR, Bex PJ, Rose DJ, Woods RL (2014) Measuring information acquisition from sensory input using automated scoring of natural-language descriptions. PLoS One 9:e93251.

Shi C, Luo G (2018) A Compact VLSI System for Bio-inspired Visual Motion Estimation. IEEE Transactions on Circuits and Systems for Video Technology 28(4):1021 – 1036.

Shioiri S, Ito S, Sakurai K, Yaguchi H (2002) Detection of relative and uniform motion. Journal of the Optical Society of America - A 19:2169–2179.

Simoncelli EP, Heeger DJ (2001) Representing retinal image speed in visual cortex. Nature Neuroscience 4:461–462.

St. Clair R, Huff M, Seiffert AE (2010) Conflicting motion information impairs multiple object tracking. Journal of Vision 10:1–13.

Tadin D, Nyquist JB, Lusk KE, Corn AL, Lappin JS (2012) Peripheral Vision of Youths with Low Vision: Motion Perception, Crowding, and Visual Search. Investigative Ophthalmology & Visual Science 53:5860–5868.

Torralba A, Oliva A (2003) Statistics of natural image categories. NETWORK: COMPUTATION IN NEURAL SYSTEMS 14:391–412.

van der Schaff A, van Hateren JH (1996) Modelling the Power Spectra of Natural Images: Statistics and Information. Vision Research 36:2759–2770.

Van Doorn A, Koenderink J (1982) Spatial properties of the visual detectability of moving spatial white noise. Experimental Brain Research 45:189–195.

Wichmann FA, Henning GB (1998) No role for motion blur in either motion detection or motion-based image segmentation. Journal of the Optical Society of America - A 15:297–306.

Wood JM, Lacherez P, Tyrrell RA (2014) Seeing pedestrians at night: effect of driver age and visual abilities. Ophthalmic and Physiological Optics 34:452–458

Wood JM, Black AA, Mallon K, Kwan AS, Owsley C (2018) Effects of Age-Related Macular Degeneration on Driving Performance. Investigative Ophthalmology & Visual Science 59:273–279.

Yang J, Stevenson SB (1997) Effects of spatial frequency, duration, and contrast on discriminating motion directions. Journal of the Optical Society of America - A 14:2041–2048.

